# Low concentrations of selenium nanoparticles enhance the performance of a generalist parasitoid and its host, with no net effect on host suppression

**DOI:** 10.1101/2023.01.12.523859

**Authors:** James Rudolph Miksanek, Charles Adarkwah, Midori Tuda

## Abstract

**BACKGROUND:** The environmental and economic costs of conventional insecticides have stirred an interest in alternative management tactics, including the use of nanotechnologies. Selenium nanoparticles (SeNPs) have many applications in agriculture but may not be compatible with biological control; however, low concentrations of SeNPs may benefit natural enemies via hormesis. This study investigates the concentration-dependent effects of SeNPs (0–1000 mg L^−1^) on *Anisopteromalus calandrae* (Howard) (Hymenoptera: Pteromalidae), a generalist parasitoid of stored product pests.

**RESULTS:** The LC_50_ of SeNPs was 1540 mg L^−1^ for female parasitoids and 1164 mg L^−1^ for males. SeNPs had a significant hormetic effect; average lifespan increased by 10% at a concentration of 4.03 mg L^−1^ for females and by 35% at 13.83 mg L^−1^ for males. In a bioassay including hosts (the azuki bean beetle, *Callosobruchus chinensis* (L.) (Coleoptera: Chrysomelidae: Bruchinae)), a low concentration of SeNPs (25 mg L^−1^) enhanced the performance of female parasitoids; lifespan increased by 23% and the number of offspring increased by 88%. However, the number of emerging hosts did not significantly decrease; in the absence of parasitism, SeNPs actually improved host emergence by 17%.

**CONCLUSION:** Because higher concentrations of SeNPs reduced parasitoid lifespan, whereas low concentrations enhanced not only parasitoid performance but also host emergence, practitioners should exercise caution when considering SeNPs for use in integrated pest management.

## 1. INTRODUCTION

The complementarity of chemical, cultural, biological, and other control strategies is a key aspect of integrated pest management (IPM)^1,2^. Because there tend to be compatibility issues between conventional insecticides and natural enemies^3–5^, and due to the environmental and economic costs of these pesticides^6,7^, there are ongoing efforts to counter an overreliance on chemical control and incorporate safer biorational alternatives to improve management outcomes and meet sustainable development goals (SDGs)^8–10^. Emerging research is also investigating the possible roles of nanotechnology in pest management, including the use of entomotoxic nanoparticles (NPs) or nano-encapsulated pesticides or botanicals^11,12–15^. Despite this effort, the ecological risks of NPs remain relatively unknown^11,16–18^. Due to the nonspecific mode of action of elemental (or elemental oxide) NPs^19^, these products might pose a risk to nontarget organisms, including natural enemies.

Even though the sublethal effects of insecticides on natural enemies are typically negative^3,20^, there are also instances in which they can be positive^21,22^. The term “hormesis” (or “hormoligosis” in insects^23^) refers to the positive or stimulatory effect of low doses of a chemical stressor, which can be characterized by a biphasic dose-response curve^24^. Since pesticide residues are often present throughout the agroecological landscape, it is important to understand how these (and other stressors) influence the biological control services provided by natural enemies^24,25^. Although there are many variables that affect the persistence of anthropogenic NPs in the environment (and their relationship to naturally occurring NPs)^18,26^, relatively small or residual amounts of NPs might offer a direct benefit to biological control agents by hormesis.

*Anisopteromalus calandrae* (Howard) (Hymenoptera: Pteromalidae) is a cosmopolitan parasitoid that attacks a broad range of hosts, including the azuki bean beetle, *Callosobruchus chinensis* (L.), and the cowpea seed beetle, *C. maculatus* (F.) (Coleoptera: Chrysomelidae: Bruchinae)^27^. As solitary idiobiont ectoparasitoids, females typically lay one egg after paralyzing a host^28^, and the developing parasitoid larva feeds from outside of its host; additional parasitoid-induced mortality can arise via host feeding^29^, a process through which adult female *A. calandrae* attack and consume the hemolymph of hosts to acquire nutrients for egg maturation^29–31^. Due to the importance of *A. calandrae* in the control of bruchine beetles and other coleopteran pests^32–36^, it is necessary to test their compatibility with the entomotoxic NPs that have garnered recent attention—selenium nanoparticles (SeNPs), in particular^37,38^—as a possible new control strategy for these insects. Furthermore, because the risks of nano-formulated insecticides are not as well studied as those of conventional insecticides^16–18^ (and more controversial^39^), these tests on *A. calandrae* may serve as an indicator of their ecotoxicity, providing insight into both the specificity of SeNPs as well as their impact on the biological control subset of ecosystem services.

SeNPs have several potential applications in agriculture: promoting plant growth, bolstering plant defenses, and acting as an antimicrobial, nematocidal, or insecticidal agent, depending on the concentration or formulation^40–42^. The entomotoxic effects of SeNPs arise through the slow release of elemental selenium, which, depending on the route of entry, may amass in the Malpighian tubules and midgut, negatively impacting the growth, development, and survival of the target insect^43–45^. While SeNPs have shown some promise as an alternative control method for bruchine beetles, including *C. chinensis*^37,38^, they may also harm the parasitoid *A. calandrae*^38^. However, there is evidence of SeNPs eliciting a hormetic response in the male hosts of *A. calandrae*^37^, so, considering the nonspecific mode of action of SeNPs, we hypothesize that a similar result will be observed for the parasitoid. The purpose of the present study is to investigate the concentration-dependent effects of SeNPs on the lifespan of the generalist parasitoid *A. calandrae* and to test the hormetic effects of a low concentration of SeNPs on parasitoid performance (lifespan, fecundity, offspring emergence time, age-dependent parasitism rate, and host suppression), all of which will provide better guidance on the utilization of SeNPs and *A. calandrae* in IPM.

## 2. METHODS

Colonies of *C. chinensis* (strain jC, originating from Japan) and its parasitoid *A. calandrae* (also from Japan) have been maintained in controlled environmental chambers under standard laboratory conditions (30°C, 60% R.H., 16:8 L:D) for more than 20 years on dried azuki beans, *Vigna angularis* var. *angularis* (Willdenow) Ohwi & Ohashi (Fabaceae) (Daiwa Grain Co., Obihiro, Japan). The *C. chinensis* colonies are maintained in large, ventilated Petri dishes (9.5 cm in diameter, 4 cm in height), whereas *A. calandrae* is maintained in standard-sized Petri dishes (10 cm in diameter, 1.5 cm in height) supplied with *C. chinensis*-infested azuki beans (16-day old pupae and late-instar larvae).

The chemical synthesis of SeNPs^46,47^ (amorphous and approximately 5–10 and 60–100 nm in diameter) was previously described by Miksanek & Tuda^37^; in brief, a 20-mL solution of SeNPs (1000 mg L^−1^) was prepared by reducing sodium selenite (43.8 mg) with ascorbic acid (2 mL, 0.633 M) in ultrapure water (17.9 mL; Milli-Q, 18.2 MΩ·cm) and stabilizing with polysorbate 20 (100 μL). Additional SeNP concentrations of 1, 10, 25, 50, 100, 200, and 500 mg L^−1^ were made by diluting the original solution with ultrapure water; a 0 mg L^−1^ control solution (containing only ultrapure water) was also prepared. All chemical reagents (sodium selenate, ascorbic acid, and polysorbate 20) were obtained from Nacalai Tesque, Inc. (Kyoto, Japan).

### 2.1 Concentration-dependent toxicity

Newly emerged (< 24 h) adult parasitoids (males and females, separately) were placed individually in mini-sized Petri dishes (35 mm in diameter, 10 mm in height). Each Petri dish has an air gap around the lid that allows for ventilation. Next, a 20-μL droplet of SeNPs (0, 1, 10, 50, 100, 200, 500, or 1000 mg L^−1^ (0–20 μg SeNPs)) was added to the center of the dish; by holding the volume constant and manipulating the concentration of SeNPs, this changes the overall dose. Each Petri dish was gently agitated to evenly cover all surfaces as well as the parasitoid with SeNPs. After this initial treatment, the parasitoids were placed in a climate-controlled chamber (30°C, 60% R.H., 16:8 L:D) and monitored daily until death. Parasitoids were not provided supplemental nutrients before, during, or after the bioassay. All eight SeNP concentrations were replicated 25 times for each sex.

Note that we refer to a “replicate” as a repetition of the complete set of treatment combinations (eight SeNP concentrations × two sexes), and, to reiterate, the parasitoids (400 in total) were subjected to treatment combinations individually rather than as part of a group. Although toxicity testing on groups of organisms is both common and convenient (with replication at the level of each set of groups), that approach is problematic, especially for *A. calandrae* because female *A. calandrae* exhibit high levels of density-dependent intraspecific interference competition^35,48,49^; if these parasitoids were exposed to SeNPs in a group, then the resulting differential rates of death across groups treated with different concentrations would introduce an increasing asymmetry in the behavioral interactions among the individuals in each group over time, which would be a confounding variable.

### 2.2 Low-concentration SeNPs and parasitoid performance

The effects of a low concentration of SeNPs on the performance of *A. calandrae*, as well as offspring emergence timeline, were tested by exposing parasitoids to infested azuki beans treated with either SeNPs or water. The purpose of this was to evaluate the complementarity of SeNPs and biological control. The experimental design consisted of a full factorial for parasitism (*A. calandrae* present or absent) and SeNP application (0 or 25 mg L^−1^ SeNPs). For this bioassay, a concentration of 25 mg L^−1^ SeNPs was chosen because a preliminary analysis of concentration-dependent survivorship suggested that this approximate concentration may best enhance parasitoid performance.

Adult parasitoids were collected from a laboratory colony within 24 h of emergence. Next, each male–female pair was treated with a 20-μL droplet of SeNPs (0.5 μg SeNPs) or ultrapure water (as previously described) and provided eight azuki beans (with a total mass of 1.1 g, on average) treated concurrently and in the same manner (by gently agitating each dish to evenly cover all surfaces as well as the beans with SeNPs) with a 30-μL droplet^37^ (0.75 μg SeNPs) (to account for the increased surface area). Each bean was infested by eight mixed-stage *C. chinensis* larvae (indirectly counted as the number of eggs on the surface of the bean; beans were visually inspected prior to use in the experiment, and those with fewer or greater than eight hatched eggs were not used); by the time of parasitoid exposure, most larvae were 15–16 days old, which is the favored host stage for parasitism; the smaller number of younger larvae are the preferred size for host-feeding. After treatment, beans and parasitoids were immediately placed in a small, untreated Petri dish (55 mm in diameter, 15 mm in height); the treated beans were not allowed to dry beforehand. After 24 h, the male parasitoid was discarded to prevent excessive sexual harassment. Parasitoid mortality was monitored at 24-h intervals. After four days, females were provided with a fresh set of infested beans (of the same treatment conditions), and the original sets of beans were checked daily to record the number and sex of emerging adults (hosts and parasitoids). The second set of beans were not monitored for daily emergence, but the total number of emerged parasitoids was recorded three weeks after the death of the parental parasitoid in order to calculate lifetime reproduction. Parasitoids were not provided supplemental nutrients before or during the bioassay. Each combination of treatments was replicated 30 times; daily emergence was only recorded from a subset of replicates (24 from each of the treatments in the absence of parasitism and 25 from each of those with a parasitoid present).

### 2.3 Modeling and Analyses

The concentration-dependent effects of SeNPs on parasitoid lifespan (age-at-death, *x*) were described using the Cedergreen–Ritz–Streibig modification of a log-logistic model, which allows for an effect of hormesis^50,51^. This model takes the form:

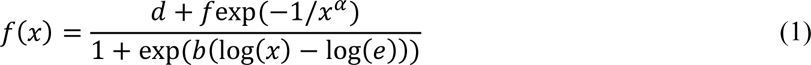

in which *d* is the upper horizontal asymptote and *f* is the magnitude of the effect of hormesis (with *α* affecting the shape of the peak); the parameters *b* and *e* have no direct biological interpretation (in the related three-parameter log-logistic model, *b* is a scaling factor and *e* is an inflection point that corresponds with the EC_50_ or LC_50_)^50,51^. If the parameter *f* is equal to 0, then there is no hormetic effect and the model simplifies to a three-parameter log-logistic curve^50,51^. Model fitting and analysis was conducted in R version 4.2.2. The term *α* was set to 0.5 when fitting the model^50,51^. Parasitoid sex was included as a grouping factor when estimating the parameters *b* and *d* (the model was simplified because preliminary analyses showed no significant effect of sex on estimates of *e* and *f*). The package *drc* was used in the analysis, which also allowed for estimating the maximum effect of hormesis from the mean response predicted by the model (using the “MAX” function^51^, as described by Cedergreen et al.^50^), lethal concentrations (e.g. LC_50_), and calculating *t*-statistics for comparisons between sexes^50,51^.

The effects of SeNPs on the lifespan and fecundity of adult female parasitoids were tested with a Cox proportional hazards model and linear regression model, respectively. A binomial GLM was used to test the effects of SeNP treatment on the sex ratio of the resulting parasitoid offspring. The effects of SeNPs on the development time of offspring (days to adult emergence) were assessed with a Cox proportional hazards model, with sex and the interaction between sex and SeNP treatment as additional factors; the identity of the maternal parasitoid was included as a random effect by clustering.

To further investigate the reproduction of female parasitoids, a linear regression was used to assess the relationship between the log-transformed early rate of reproduction (the age-corrected number of offspring per day, calculated as the number of offspring produced from days 0–3 (inclusive) divided by the lifespan of the parasitoid) and SeNP treatment on the reproductive rate later in life (from day 4–death, calculated in the same fashion); only parasitoids with a lifespan greater than four days and a nonzero early reproductive rate were included in this analysis. Next, to investigate the influence of SeNPs on the lifetime reproductive success (total number of offspring) of female parasitoids as a function of age, an asymptotic Michaelis–Menten-type model was fit to the data. This model predicts the total number of offspring of a female parasitoid living to age *t* as:

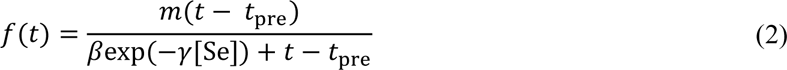

Analogous to the standard Michaelis-Menten model, the parameter *m* denotes the upper asymptote of the saturating curve and *β* is associated with the midpoint of the saturating curve; biologically, *m* represents the maximum number of parasitoid offspring (limited by host depletion or as a more general effect of parasitoid senescence). The modifications on the Michaelis-Menten model include an *x*-intercept (*t*_pre_) and the effect of SeNPs (exp(*−γ*[Se])). The term *t*_pre_ applies a horizontal transformation to the curve; biologically, *t*pre represents the length of the pre-oviposition period of the adult female (a female might use this time for mating, sperm storage, host feeding, or egg maturation). The function exp(*−γ*[Se]) applies an effect *γ* of SeNPs ([Se] = 0 or 25 mg L^−1^) to *β*; thus, at a concentration of 0 mg L^−1^ (or if there is no effect *γ*), then *β* is unaffected. For simplicity, the Michaelis–Menten-type model only considers the effect of SeNPs on the parameter *β*; however, this formulation was also supported by an informal analysis of the shortest- and longest-lived parasitoids (0–3 and 9–12 days, respectively), in which were there no differences in the reproduction of control and SeNP-treated parasitoids in either age class, suggesting that *m* and *t*_pre_ are likely unaffected by SeNP treatment.

Finally, the effects of SeNPs and parasitism on the number and sex ratio of emerging hosts were tested with a linear mixed-effects model (LMM) and binomial generalized mixed-effects model (GLMM), respectively. In both models, the identity of the source colony was included as a random effect to account for potential variation in the stage structure or exact number of hosts; because source colony did not improve the statistical models of parasitoid life history (described above), this random effect was only included in these analyses of the host population.

All modeling and statistical analyses were conducted in R version 4.2.2. The Michaelis–Menten-type model was fit using nonlinear least squares; confidence intervals for model predictions were calculated by Monte Carlo simulation (*n* = 100000). The packages *car*, *lme4*, *merTools*, *survival*, *coxed*, and *propagate* were also used in the analyses.

## 3. RESULTS

### 3.1 Concentration-dependent toxicity

The lifespan of adult parasitoids was measured in response to a range of SeNP concentrations (0–1000 mg L^−1^). In fitting the Cedergreen–Ritz–Streibig-modified log-logistic model to the concentration-dependent effects of SeNPs on the lifespan of male (*n* = 200) and female (*n* = 200) parasitoids (with parasitoid sex as a grouping factor when estimating *b* and *d*), the estimates for all parameters were significant (Figure 1, Table 1). The upper asymptote *d*—which represents the maximum lifespan in the absence of SeNPs—was 4.55 ± 0.15 d (estimate ± SE) for females and 3.01 ± 0.16 d for males, and the difference between sexes was significant (*t* = 6.27, *p* < 0.001) (Figure 1; Table 1); the estimates of *b* (a scaling factor) were marginally different between sexes (*t* = −1.91, *p* = 0.057) (Table 1).

**Figure 1.**
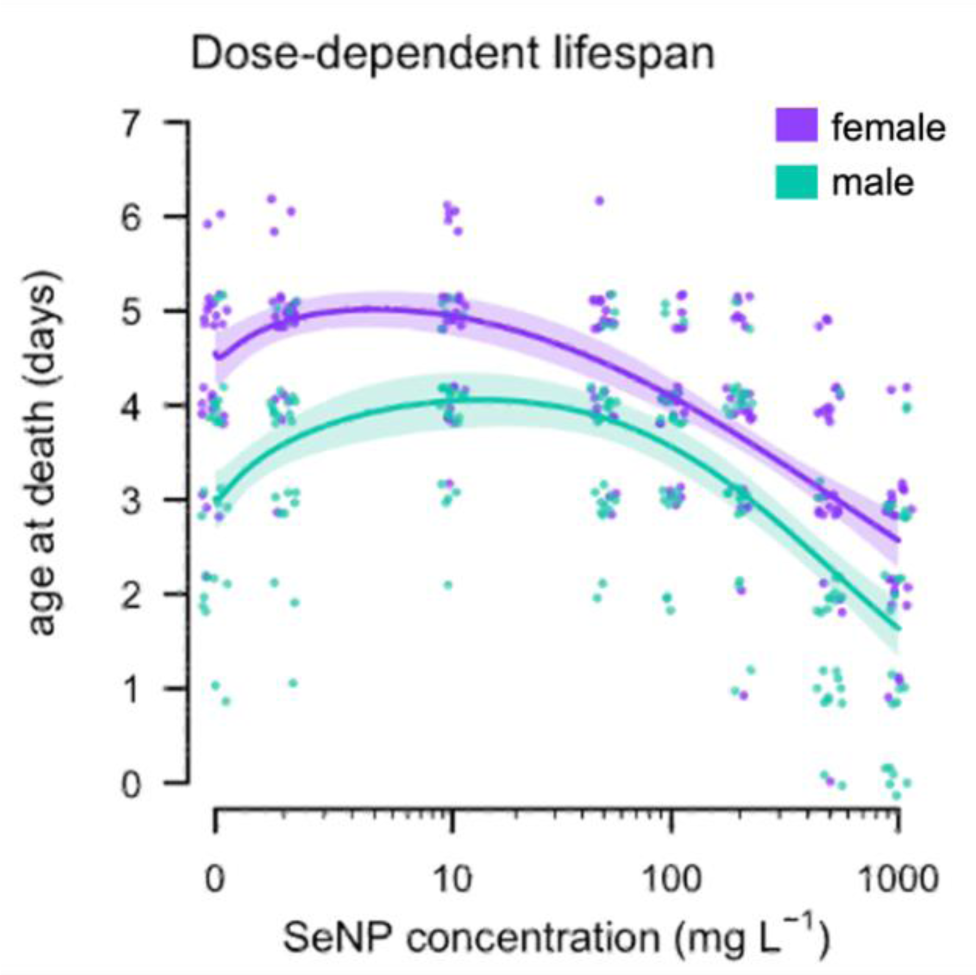
Concentration-dependent lifespan of the parasitoid *Anisopteromalus calandrae* (adult lifespan, i.e. age at death) treated individually with different concentrations of selenium nanoparticles (SeNPs) in the absence of hosts; the solid line plots the Cedergreen–Ritz–Streibig-modified log-logistic model (with 95% CI); points plot observed values (jittered for clarity).

**Table 1.** Parameter estimates for the Cedergreen–Ritz–Streibig-modified log-logistic model for the concentration-dependent effect of selenium nanoparticles (0–1000 mg L^−1^) on the lifespan of the parasitoid *Anisopteromalus calandrae* in the absence of hosts.

| Parameter | Grouping (sex) | Estimate | SE | *t*-value | *p*-value |
| --- | --- | --- | --- | --- | --- |
| *b* | Female | 0.47 | 0.09 | 5.29 | < 0.001 |
|  | Male | 0.84 | 0.16 | 5.24 | < 0.001 |
| *d* (upper asymptote) | Female | 4.55 | 0.15 | 29.60 | < 0.001 |
|  | Male | 3.01 | 0.16 | 19.21 | < 0.001 |
| *e* | – | 494.80 | 118.95 | 4.16 | < 0.001 |
| *f* (effect of hormesis) | – | 1.63 | 0.44 | 3.73 | < 0.001 |

The LC_50_ was 1540 ± 391 mg L^−1^ for females (estimate ± SE) and 1164 ± 224 mg L^−1^ for males, but there was no significant difference in the estimates of LC_50_ between sexes (*t* = 1.19, *p* = 0.237). For hormetic effects, the estimated age at death of SeNP-treated females was 5.02 d at a concentration of 4.03 mg L^−1^; and the estimated age at death of SeNP-treated males was 4.06 d at a concentration of 13.83 mg L^−1^ (Figure 1; Table 1).

### 3.2 Low-concentration SeNPs and parasitoid performance

Parasitoid performance was assessed in response to a low concentration (25 mg L^−1^) of SeNPs to investigate the effects of hormesis on a generalist biological control agent. Adult female parasitoids (*n* = 60) treated with 25 mg L^−1^ SeNPs lived significantly longer than control parasitoids; adult lifespan was 7.53 ± 0.52 and 6.10 ± 0.48 d, respectively (mean ± SE) (Figure 2a; Table 2). SeNP-treated females also produced a greater number of offspring; the total number of offspring was 13.0 ± 1.1 vs. 6.9 ± 1.1 (Figure 2b; Table 2). The sex ratio (*n* = 56) and development time (*n* = 285) of the offspring of SeNP-treated parasitoids did not differ from those of the control (Table 2). Regardless of SeNP treatment, female offspring (*n* = 233) emerged later than males (*n* = 52); the time to emergence was 13.66 ± 0.09 and 12.73 ± 0.18 d, respectively (Table 2).

**Figure 2.**
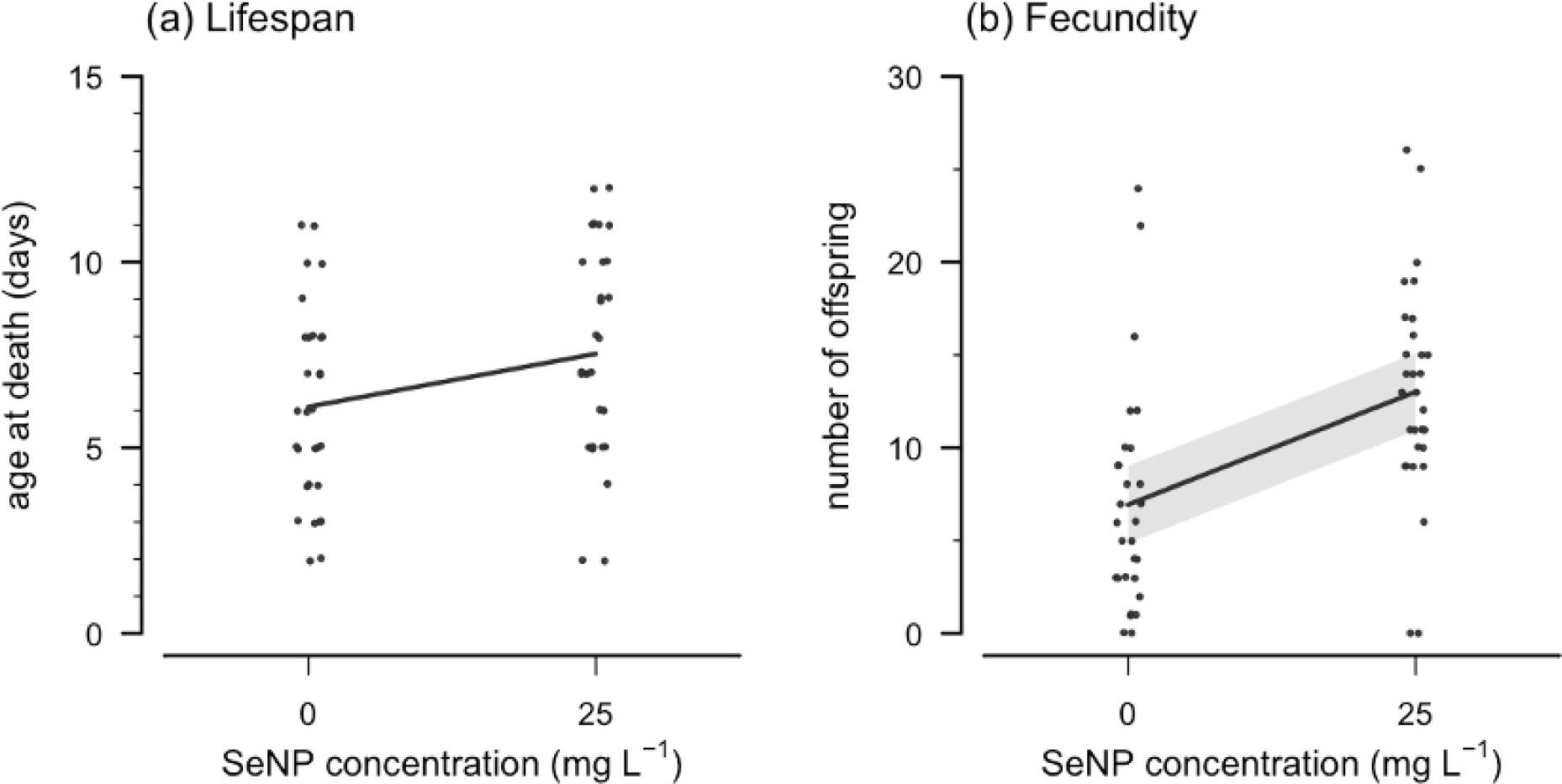
Lifespan and fecundity of adult female *Anisopteromalus calandrae* as a function of selenium nanoparticle (SeNP) treatment (0 or 25 mg L^−1^). **(a)** Lifespan of adult female parasitoids; the solid line indicates the average age at death. **(b)** Fecundity (total number of offspring); the solid line plots the linear regression model (with 95% CI). In both panels, points plot observed values (jittered for clarity).

**Table 2.** Statistical tests of the effects of selenium nanoparticles (25 mg L^−1^) on the life history parameters of adult female parasitoids (*Anisopteromalus calandrae*) and offspring.

| Response (Model) | Predictor | df | $\chi^2$ | *p*-value |
| --- | --- | --- | --- | --- |
| Age at death (Cox prop. hazards) | SeNP treatment | 1 | 4.27 | 0.039 |
| Reproduction (linear regression) | SeNP treatment | 1 | 16.24 | < 0.001 |
| Sex ratio (binomial GLM) | SeNP treatment | 1 | 0.89 | 0.346 |
| Development time (Cox prop. hazards) | Sex | 1 | 18.38 | < 0.001 |
|  | SeNP treatment | 1 | 0.03 | 0.856 |
|  | Sex × SeNP treatment | 1 | 0.02 | 0.884 |

For ovipositing female parasitoids with an age of death > 4 d (*n* = 47), the early reproductive rate was inversely correlated with the rate of reproduction later in life (offspring per day, days 4–death) (*F*_3,43_ = 14.94, multiple *R*^2^ = 0.510, *p* < 0.001) (Figure 3a). As the early rate of reproduction increased, there was a decrease in the rate of reproduction later in life; treatment with 25 mg L^−1^ SeNPs increased the rate of reproduction overall (Figure 3a; Table 3a). To investigate the lifetime reproductive success of all female parasitoids (*n* = 60), a Michaelis–Menten-type model was implemented, in which all parameter estimates were significant (Figure 3b; Table 3b). The maximum number of offspring *m* was 44.7 ± 17.8 (estimate ± SE) and the preoviposition period was 1.90 ± 0.30 d (Table 3b). There was also a significant effect of SeNP treatment *γ*, with 25 mg L^−1^ SeNPs increasing the lifetime reproductive success of female parasitoids (Figures 3a and 3b; Tables 3a and 3b).

**Figure 3.**
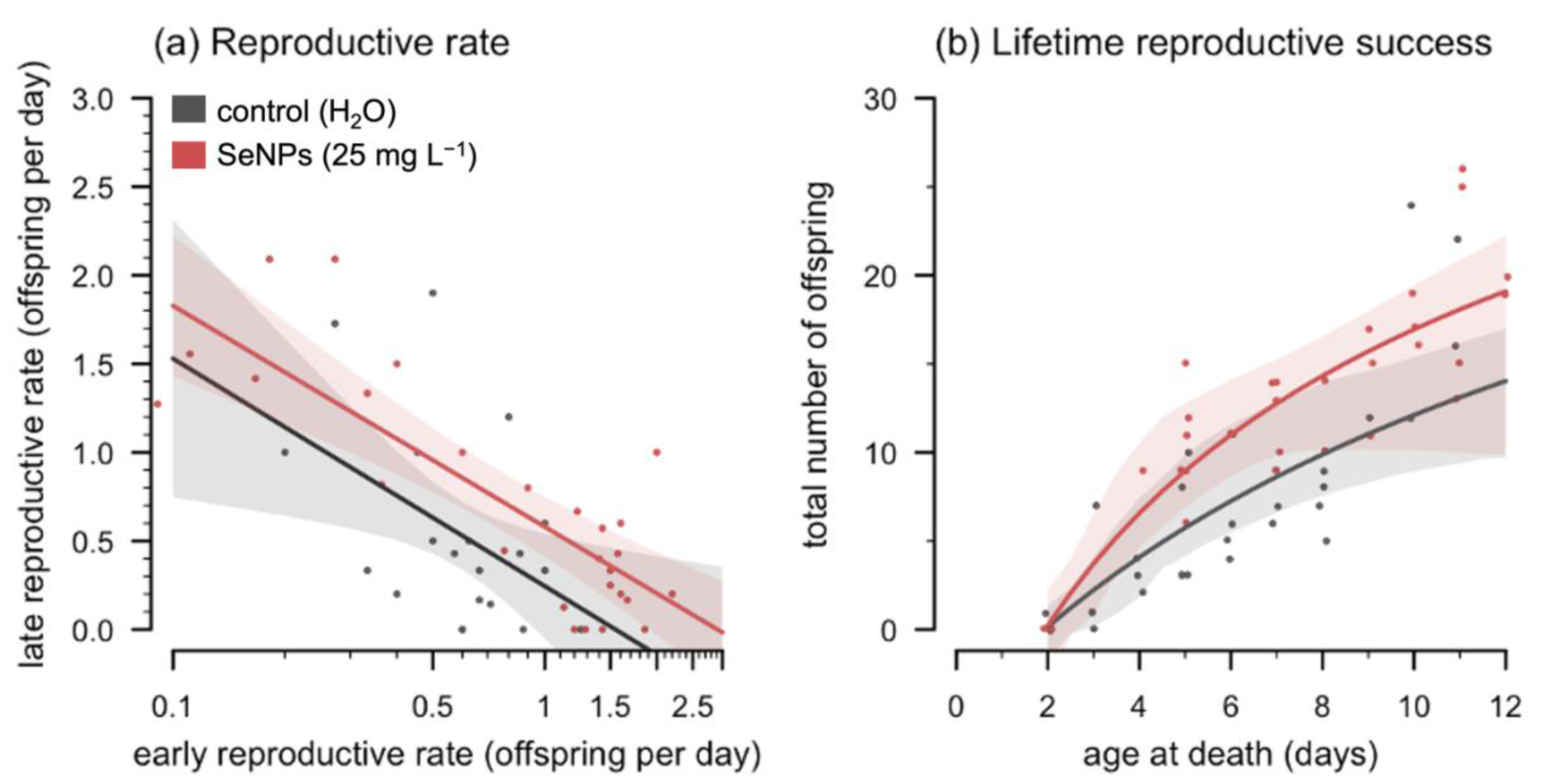
Reproduction of the generalist parasitoid *Anisopteromalus calandrae* as a function of age and selenium nanoparticle (SeNP) treatment (0 or 25 mg L^−1^). **(a)** Correlation between the early and late reproductive rates of female parasitoids (number of offspring per day [age-adjusted], days 0–3 vs. day 4– death); the solid line plots the linear regression model (with 95% CI). **(b)** Lifetime reproduction, or the total number of offspring, per female; solid lines plot the Michaelis-Menten-type models (with 95% CI). In both panels, points plot observed values (jittered for clarity in panel b).

**Table 3a.** Correlation between early and late reproduction (number of offspring per day) by female *Anisopteromalus calandrae* and the effects of selenium nanoparticles (25 mg L^−1^).

| Predictor | df | $\chi^2$ | *p*-value |
| --- | --- | --- | --- |
| Early reproductive rate<br>[log(age-adjusted number of offspring per day)] | 1 | 42.97 | < 0.001 |
| SeNP treatment | 1 | 6.32 | 0.016 |
| Early reproductive rate × SeNP treatment | 1 | 0.00 | 0.946 |

**Table 3b.** Parameter estimates for the Michaelis–Menten-type model of the lifetime reproductive success of selenium nanoparticle-treated (25 mg L^−1^) female parasitoids (*Anisopteromalus calandrae*).

| Parameter | Estimate | SE | *t*-value | *p*-value |
| --- | --- | --- | --- | --- |
| Maximum number of parasitized hosts, *m* | 44.72 | 17.85 | 2.51 | < 0.001 |
| $\beta$ | 21.94 | 11.91 | 1.84 | < 0.001 |
| Preoviposition period, *t*<sub>pre</sub> | 1.90 | 0.30 | 6.35 | < 0.001 |
| Effect of SeNP treatment, $\gamma$ | 0.02 | 0.01 | 3.62 | < 0.001 |

Finally, the effects of SeNPs and parasitism on the host population were evaluated as the number and sex ratio of the emerging hosts in each Petri dish (*n* = 120). The number of emerging hosts was lower in the presence of the parasitoid than in the absence of the parasitoid but, when treated with SeNPs, was higher in the absence of parasitism (Figure 4a, Table 4). When treated with 0 mg L^−1^ SeNPs, there were an estimated 21.5 (95% prediction interval: 15.0–28.8) emerging hosts per Petri dish in the absence of parasitism and 19.1 (12.3–26.1) emerging hosts per dish in the presence of a female parasitoid; when treated with 25 mg L^−1^ SeNPs, there were 25.1 (18.6–31.4) emerging hosts per dish in the absence of parasitism and 17.1 (10.9–23.4) emerging hosts with a parasitoid present (Figure 4a, Table 4). Finally, the sex ratio of the emerging hosts shifted from a male bias (0.54 [0.51–0.57]) at 0 mg L^−1^ SeNPs to a slight female bias (0.47 [0.45–0.51]) when treated with 25 mg L^−1^ SeNPs, independent of parasitoid presence (Figure 4b, Table 4).

**Figure 4.**
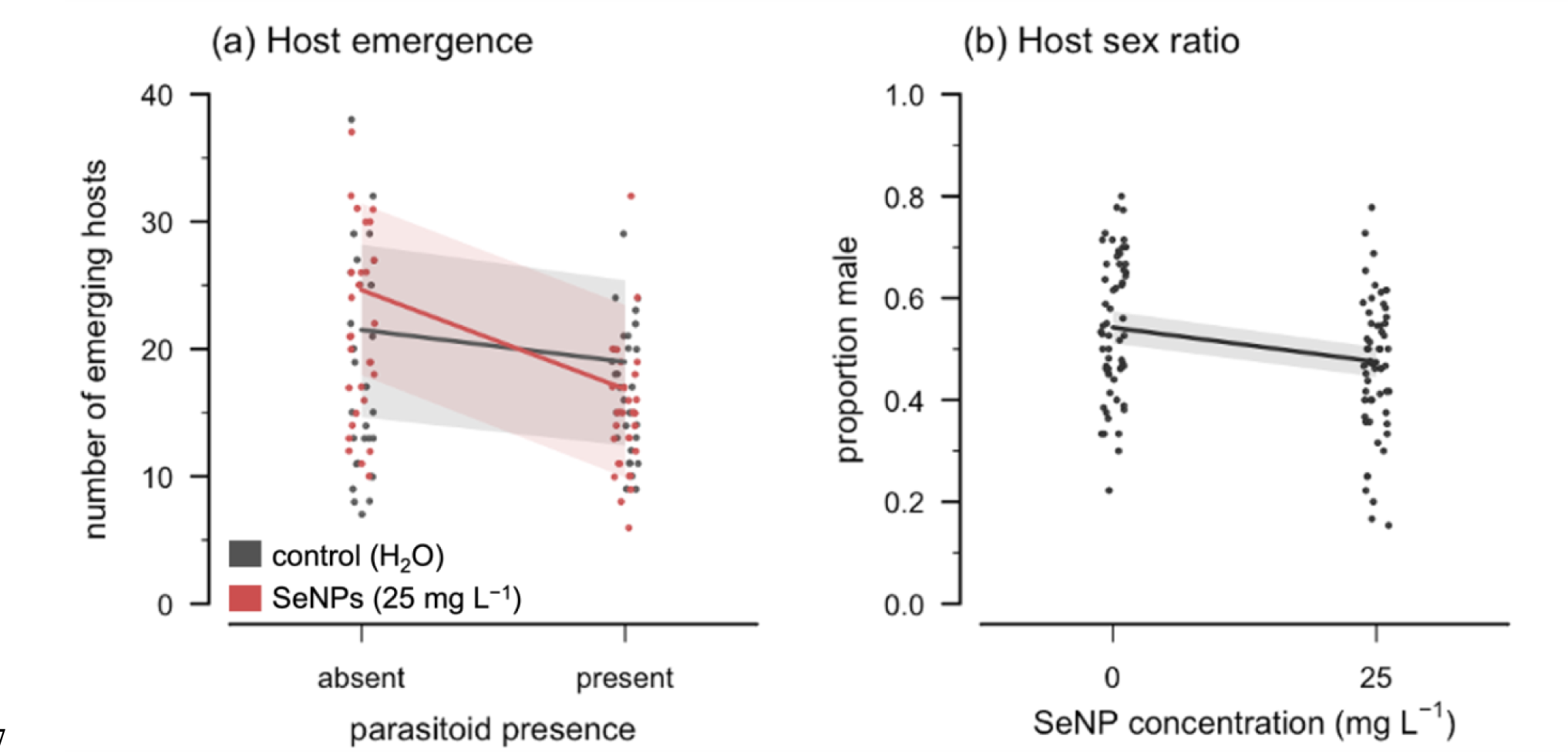
Influence of parasitism and selenium nanoparticle (SeNP) treatment (0 or 25 mg L^−1^) on the host/pest population. **(a)** Escaped hosts, or the number of emerging adult *Callosobruchus chinensis*; solid lines plot the population-level predictions from the LMM (with 95% prediction interval). **(b)** Sex ratio of emerging beetles; the solid line plots the population-level predictions from the binomial GLMM (with 95% prediction interval). In both panels, points plot observed values (jittered for clarity).

**Table 4.** Statistical tests of the effects of the parasitoid *Anisopteromalus calandrae* and selenium nanoparticles (25 mg L^−1^) on the host/pest (*Callosobruchus chinensis*) population.

| Response (Model) | Predictor | df | $\chi^2$ | *p*-value |
| --- | --- | --- | --- | --- |
| Host emergence<br>(LMM) | Parasitoid presence | 1 | 26.91 | < 0.001 |
|  | SeNP treatment | 1 | 0.61 | 0.436 |
|  | Parasitoid presence × SeNP treatment | 1 | 8.14 | 0.004 |
| Host sex ratio<br>(binomial GLMM) | Parasitoid presence | 1 | 0.06 | 0.808 |
|  | SeNP treatment | 1 | 9.22 | 0.002 |
|  | Parasitoid presence × SeNP treatment | 1 | 0.13 | 0.715 |

## 4. DISCUSSION

Because of the environmental and economic costs of conventional chemical insecticides, there is ongoing interest in exploring alternative management tactics, including the use of nanotechnologies. Selenium nanoparticles (SeNPs) have many applications in agriculture, but their nonspecific entomotoxic mode of action may limit their compatibility with biological control agents; however, low concentrations may benefit natural enemies via hormesis. The present study found that, for the generalist parasitoid *A. calandrae*, the LC_50_ of SeNPs was 1540 mg L^−1^ for female parasitoids and 1164 mg L^−1^ for males, and an analysis of the concentration-response curves identified a significant hormetic effect; for females, the estimated age at death increased by 10% at a concentration of 4.03 mg L^−1^ and, for males, increased by 35% at 13.83 mg L^−1^. In a follow-up bioassay that provided female parasitoids with hosts, the low concentration of SeNPs was found to enhance parasitoid performance; female lifespan increased by 23% and the number of offspring increased by 88%. Regardless, the number of emerging hosts (*C. chinensis*) did not significantly decrease after treatment with low-concentration SeNPs but rather, in the absence of parasitism, increased by 17%. This indicates that, while a low concentration of SeNPs did enhance parasitoid performance, there was no net effect on host suppression because *C. chinensis* responded positively in parallel.

There exists a delicate balance in the essential role of selenium as a micronutrient and its dose-dependent toxicity in both vertebrate and invertebrate animals^40–45,52–56^. In insects, diets with supplemental selenium can benefit growth, survival, and reproduction, but higher concentrations can have the opposite effect^40,43–45,53^. While insects play a role in the biogeochemical cycling of selenium^54,56^, little is known of the impact of selenium on predator–prey or host–parasitoid ecology; the biotransfer or biomagnification^57^ of selenium can negatively affect the development and fitness of predators^58^, and higher concentrations of selenium can potentially disrupt host–parasitoid population dynamics, with *Cotesia marginiventris* (Cresson) (Hymenoptera: Braconidae) and the beet armyworm, *Spodoptera exigua* (Hubner) (Lepidoptera: Noctuidae), serving as an example^56,59^. Even though SeNPs have a lower toxicity and higher bioavailability than inorganic or organic forms of selenium^42^, applications of SeNPs ranging in concentration from 100–500 mg L^−1^ can still have significant entomotoxic effects^42,60^, including on *C. chinensis*^37,61^. However, in *C. chinensis*, endosymbiotic bacteria can mediate resistance to SeNPs^37^; *Wolbachia* provides females with resistance to low concentrations of SeNPs, whereas higher concentrations had an entomotoxic (but not insecticidal) effect regardless of infection status, decreasing life expectancy and reducing fecundity^37^.

In the present study of the *C. chinensis*–*A. calandrae* system, the parasitoid *A. calandrae* also began showing adverse effects within the 100–500 mg L^−1^ range. Thus, the concurrent use of SeNPs and biological control would not be compatible in this case. Similarly, since the pest species also appears to exhibit a hormetic response to a low concentration of SeNPs, this limits the complementary of SeNPs and biological control, at least in the short term; additional studies that extend beyond the single-parasitoid, single-generation paradigm (and that also address the timing of SeNP applications and residual effects), would be required to identify any long-term consequences that might be relevant for pest suppression. But first, there is the fundamental question of whether the performance of *A. calandrae* is enhanced as a result of a response to SeNPs as a chemical stressor (i.e. hormoligosis/hormesis), via the potential benefits of selenium as a micronutrient (parasitoids could be observed drinking from dispersed droplets of SeNPs [or water] before the application completely dried, although this would not offer the same benefits as supplemental honey or sugar^30,62^), or through some combination of the two.

The positive effects of a low concentration of SeNPs on lifespan (and fecundity) were more prominent when hosts were available for host feeding and oviposition, which suggests that SeNPs may at least alter the nutritional physiology of female parasitoids (and, if linked to host feeding, this may introduce an additional source of density dependence in the system^63,64^). However, because selenoproteins have largely been lost in the related parasitoid *Nasonia vitripennis* (Walker) (Hymenoptera: Pteromalidae) and the honey bee *Apis mellifera* L. (Hymenoptera: Apidae)^65,66^, the positive effects of low-concentration SeNPs are more likely to be a hormetic response to small amounts of oxidative stress or via a reduced stress threshold rather than as a positive response to an essential micronutrient^67–69^. Regardless, because most Coleoptera also lack selenoproteins^65,66,69^, a hymenopteran parasitoid and its coleopteran host might be expected to exhibit similar physiological responses to SeNPs, which could complicate control efforts.

## 5. CONCLUSION

Because higher concentrations of SeNPs reduced parasitoid lifespan, the concurrent use of SeNPs and biological control do not appear to be compatible tactics in the IPM of *C. chinensis*. This is an important finding because it offers insight into the potential ecotoxicity of SeNPs and it also raises concerns of the nonspecific mode of action of SeNPs, which would be antithetical to their purpose as an alternative to conventional chemical controls. Additionally, because a low-concentration treatment of SeNPs benefitted not only the parasitoid but the host as well, SeNPs and biological control may not be complementary tactics, either. Future studies on the residual effects of SeNP applications, on the trade-offs between parasitoid lifespan and reproduction, or on population-level responses (extending beyond single-parasitoid, single-generation experiments) might offer additional insight. It remains possible that, within some broader context, SeNPs might still have a role in IPM or biological control.

## 6. ACKNOWLEDGEMENTS

## 6.1 Author contributions

All authors contributed to the study conception and design. Material preparation and data collection were performed by JRM and CA. JRM and MT participated in data analysis. The first draft of the manuscript was written by JRM, and all authors commented on subsequent versions of the manuscript. All authors read and approved the final manuscript.

## 6.2 Data availability

The datasets and R script associated with the current study are available from the primary corresponding author on request.

## 6.3 Funding

JRM is an International Research Fellow of Japan Society for the Promotion of Science (JSPS). CA was supported by an Invitational Fellowship from JSPS. MT received additional support by KAKENHI (19K06840) from JSPS.

## 7. CONFLICT OF INTEREST STATEMENT

The authors declare no conflicts of interest and affirm that the present study was carried out in adherence to all applicable ethical standards.

## Notes

### Competing Interest Statement

The authors have declared no competing interest.

### Summary of Updates

Revisions were made in response to the detailed and thoughtful comments of two expert reviewers.

